# Withaferin A inhibits LFA-1-stimulated ZAP70 activity and T-cell motility

**DOI:** 10.1101/2021.04.25.441369

**Authors:** Mobashar Hussain Urf Turabe Fazil, Chandra Sekhar Chirumamilla, Claudina Perez-Novo, Sunil Kumar, Siu Kwan Sze, Wim Vanden Berghe, Navin Kumar Verma

**Affiliations:** Lee Kong Chian School of Medicine, Nanyang Technological University Singapore, Clinical Sciences Building, 11 Mandalay Road, Singapore; Laboratory of Protein Chemistry, Proteomics and Epigenetic Signaling (PPES), Department of Biomedical Sciences, University of Antwerp, Belgium; Indian Council of Agricultural Research-National Bureau of Agriculturally Important Microorganisms, Kushmaur, Mau, Uttar Pradesh, India; School of Biological Sciences, Nanyang Technological University Singapore, Singapore

**Keywords:** T-lymphocytes, migration, withaferin A, inflammation, kinome

## Abstract

Here we report that a steroidal lactone withaferin A (WFA) can inhibit T-cell motility, which is crucial for adaptive immune responses as well as autoimmune reactions. Tandem mass spectrometry identified WFA-interactome in human T-cells that were stimulated to migrate *via* cross-linking of the lymphocyte function-associated antigen-1 (LFA-1) integrin with the ligand intercellular adhesion receptor 1 (ICAM-1). Data revealed significant enrichment of the zeta-chain-associated protein kinase 70 (ZAP70) and cytoskeletal actin protein interaction networks. Phospho-peptide mapping and kinome analysis substantiated kinase signaling downstream of ZAP70 and cytoskeletal kinase pathways as key WFA targets, which was further confirmed by *in silico* analysis and molecular assays. The WFA-ZAP70 complex was disrupted by a redox agent dithiothreitol, suggesting a covalent binding interface. Moreover, WFA ablated the phosphorylation of the myosin light chain, further constraining T-cell motility. These studies identify a mechanism whereby WFA can impact T-cell motility. WFA can therefore be exploited to pharmacologically controlling host immune responses and preventing autoimmune-mediated pathologies.

---

T-cell motility is an intrinsic attribute that ensures immuno-surveillance as well as mounting of an adaptive immune response (1). At the local site of inflammation, T-cells migrate to facilitate homeostasis as well as resolve tissue damage by a coordinated array of regulated signal transduction and cytoskeletal remodelling mechanisms (2). A critical factor in the regulation of T-cell motility is the adhesive engagement of the T-cell αLβ2 integrin lymphocyte function-associated antigen-1 (LFA-1) with the ligand intercellular adhesion receptor 1 (ICAM-1), expressed on the endothelium within the inflamed tissue (3,4). The interaction between LFA-1 and ICAM-1, activates an array of kinases triggering downstream signaling cascades that facilitate cytoskeletal remodelling and transendothelial migration of T-cells to tissue sites (4–7).

Withaferin A (WFA), an archetype withanolide, discovered from the root extract of *Withania somnifera*, is well known for immunomodulatory properties (8). Little is known about the effect of this compound on T-lymphocyte migration.

Previous studies using various cell types have shown a broad range of WFA targets, including kinases like I-kappa-B-kinase β (IKKβ) and p38 mitogen-activated protein (MAP) kinase (9). However, an integrated kinome-wide response and the molecular basis for such biological activities of WFA on T-cell motility are unclear.

In the current study, we determined intracellular effects of WFA on the migratory behaviour of human T-lymphocytes. We performed a phosphopeptidome analysis using phospho-peptide substrate microarrays and identified kinase targets of WFA in LFA-1-stimulated T-cells. Proteomics, *in silico* analysis and molecular assays confirmed WFA-interaction partners in motile T-cells. We demonstrate that WFA inhibits T-cell motility by a mechanism involving an interplay between signaling cascades downstream of the zeta-chain-associated protein kinase 70 (ZAP70) and the actin cytoskeleton.

## Results

### WFA inhibits LFA-1-stimulated T-cell motility without impacting cell adhesion on ICAM-1

Immediately after engagement of the LFA-1 receptor, T-cells polarize and undergo dynamic cytoskeletal remodelling exhibiting a tadpole-like migratory phenotype. To determine the effect of WFA on T-cell motility, we pre-treated human primary T-cells with increasing concentrations of WFA (0.3 to 2.5 μM) for 3 h and then seeded onto recombinant ICAM-1 (rICAM-1)-coated wells of 96-well plates. Motile T-cells acquired a distinct polarized and elongated morphology. However, while WFA-treated cells adhered to the rICAM-1-coated surface, they displayed loss of migratory morphologies (Fig. 1*A*). Multiparametric quantitation of migratory phenotypes employing high content analysis showed that WFA dose-dependently decreased cell 1/form factor (a measure of cell polarity), cell area and nuclear displacement; 1.25 μM WFA completely inhibited T-cell migration without impacting the number of cells adhered to immobilized rICAM-1 (Fig. 1*B*) and cell viability (Fig. 1*C* and S1). Therefore, this concentration of WFA (1.25 μM) was chosen for further analysis.

**Figure 1.**
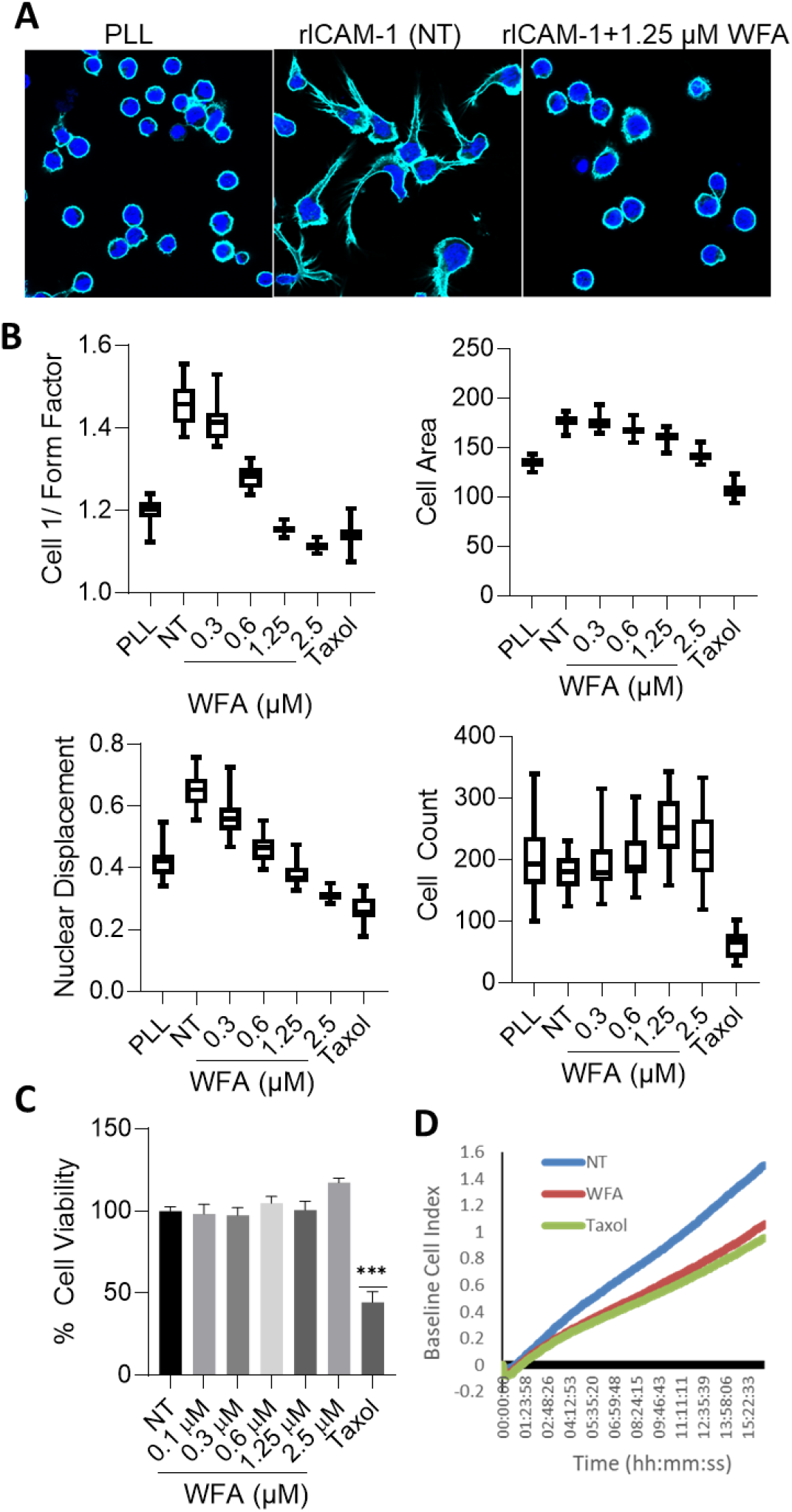
WFA inhibits T-cell migration. Human primary T-cells were pre-treated with increasing concentrations of WFA (0.1 to 2.5 µM) for 3 h or left untreated (*NT*, treated with 0.001% DMSO as a control). Cells were allowed to migrate on immobilized rICAM-1. Resting cells were placed on poly-L lysine (*PLL*)-coated plate. (**A**) Cells were fixed after 30 min of migration, stained with Hoechst (*blue*) to visualize nuclei and Alexa Fluor® phalloidin 647 (*cyan*) to visualize cell periphery/actin cytoskeleton and imaged by an automated microscopy. (**B**) Box plots showing interquartile range with median values recorded in 16 random fields from high content analysis - evaluated for cell 1/form factor, cell area, nuclear displacement, and cell count/field. (**C**) Human primary T-cells (2×10^4^ cells/per well in 96-well plates in triplicates) were treated with increasing concentrations of WFA for 3 h, viability of cells was determined using an MTS-based assay and absorbance readings normalized against untreated (*NT*) controls is plotted as % cell viability (mean ± SD, 3 independent experiments performed in triplicates); ***, p<0.0001. (**D**) Realtime chemotaxis determination by impedance-based measurements of WFA-treated (1.25 μM) human primary T-cells towards the chemokine SDF-1α. Untreated (NT) migrating cells and cells treated with taxol were used as controls. Baseline was drawn automatically for wells without SDF-1α. Data is representative of three independent experiments.

We next used a trans-well assay to assess the effect of WFA on T-cell chemotaxis towards the chemokine stromal cell-derived factor 1 α (SDF-1α). Human primary T-cells pre-treated with 1.25 μM WFA showed significantly reduced chemotaxis in comparison to the control, as quantified by impedance-based measurements in real-time (Fig. 1*D*). Notably, WFA-treated human primary T-cells were able to adhere on the rICAM-1-coated surface under flow in the same way as untreated cells (Movies S1 and S2). These results indicate that WFA inhibits LFA-1-induced T-cell motility and chemotaxis by affecting downstream signalling cascades.

### WFA interactome in motile T-cells

To identify WFA target intracellular molecules (WFA-interactome) in motile T-cells, we pulled-down cellular proteins using WFA-Biotin and streptavidin beads and performed mass spectrometry analysis. A schematic of the proteomics workflow is depicted in Fig. 2. A total of 309 unique proteins were identified using a peptide threshold of 99%, minimum 2 peptide cut-off and a false discovery rate ≤ 1.9% (Table S1). In particular, we identified several kinases in the T-cell WFA-interactome, including ZAP70, IKKβ, STK10 and pyruvate kinase, in addition to mitochondrial enzymes (methylcrotonoyl-coA carboxylase 1, citrate synthase, glutamate dehydrogenase), cytoskeletal proteins (filamin, myosin, vimentin, keratin, actin, tubulin), glucose 6 phosphate dehydrogenase, Ras-GTPase activating protein and fatty acid synthase. Ingenuity Pathway Analysis® (IPA®) revealed 3 central nodes with ZAP70, nuclear factor kappa B (NFκB) and STAT1 proteins interconnected *via* the TNFα pathway in WFA-treated LFA-1-stimulated T-cells (Fig. 2).

**Figure 2.**
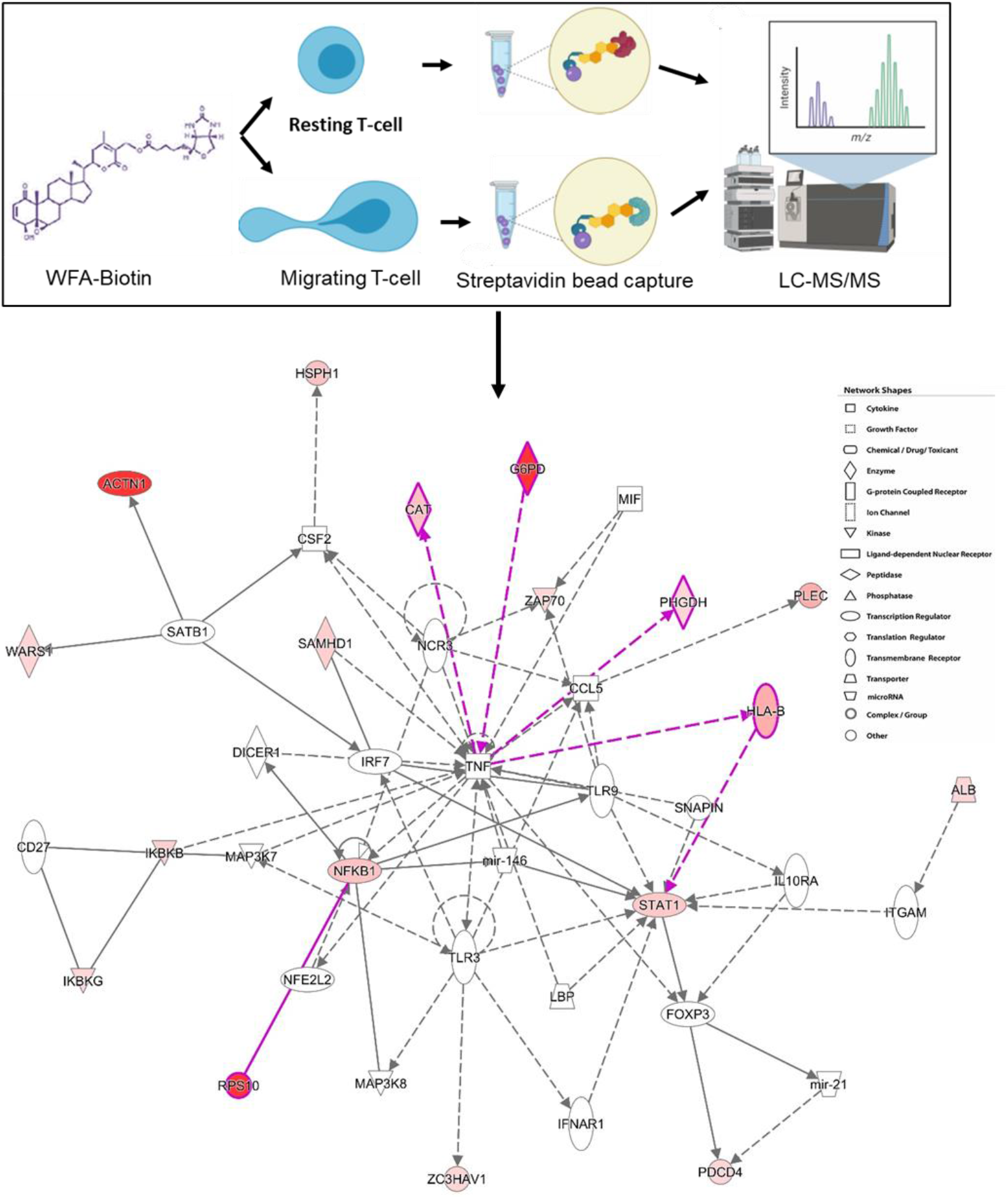
Identification of WFA interactome in migrating T-cells. Schematic representation of workflow employed in mass spectrometry-based proteomics analysis of WFA interactome in human T-cells. An image output from IPA® indicating statistically probable interactions of WFA in LFA-1-stimulated T-cells is shown. Proteins depicted in red circles are high abundance in WFA treated migrating T-cells, compared to resting T-cells. The type of network, shapes in IPA® are provided in the legend.

### WFA targets cellular kinases involved in T-cell motility

A preliminary comparison of resting and migrating T-cells indicated a significant change in the overall kinome profile of LFA-1-stimulated migrating T-cells (Fig. 3). Average phosphopeptide intensity calculations of the peptides representing KCNA2/3, FRAP, TAU, and RB1 within the serine/threonine kinase (STK) group and PGFRB and CD3ζ within the protein tyrosine kinase (PTK) family were repressed in motile T-cells (Fig. 3). Reciprocally, increased intensities were observed for DYRK1A, CRK, EPHB1, MAPK10, glycogen phosphorylase, phospholemman and CFTR specific phosphopeptides in migrating T-cells as compared to resting T-cells. Although ZAP70 peptide phosphorylation on the array was increased in LFA-1-stimulated motile T-cells, upstream kinase analysis of the total phosphopeptidome linked to ZAP70 within the array showed an overall diminution in related-kinases activity in motile T-cells (Fig. 4). This suggests an ongoing dynamic kinases/phosphatases interplay with function of time in migrating T-cells. This may also be due to loss of information regarding activity in upstream kinase analysis, as multiple peptide spots may correspond to unique sites on a specific protein in certain cases.

**Figure 3.**
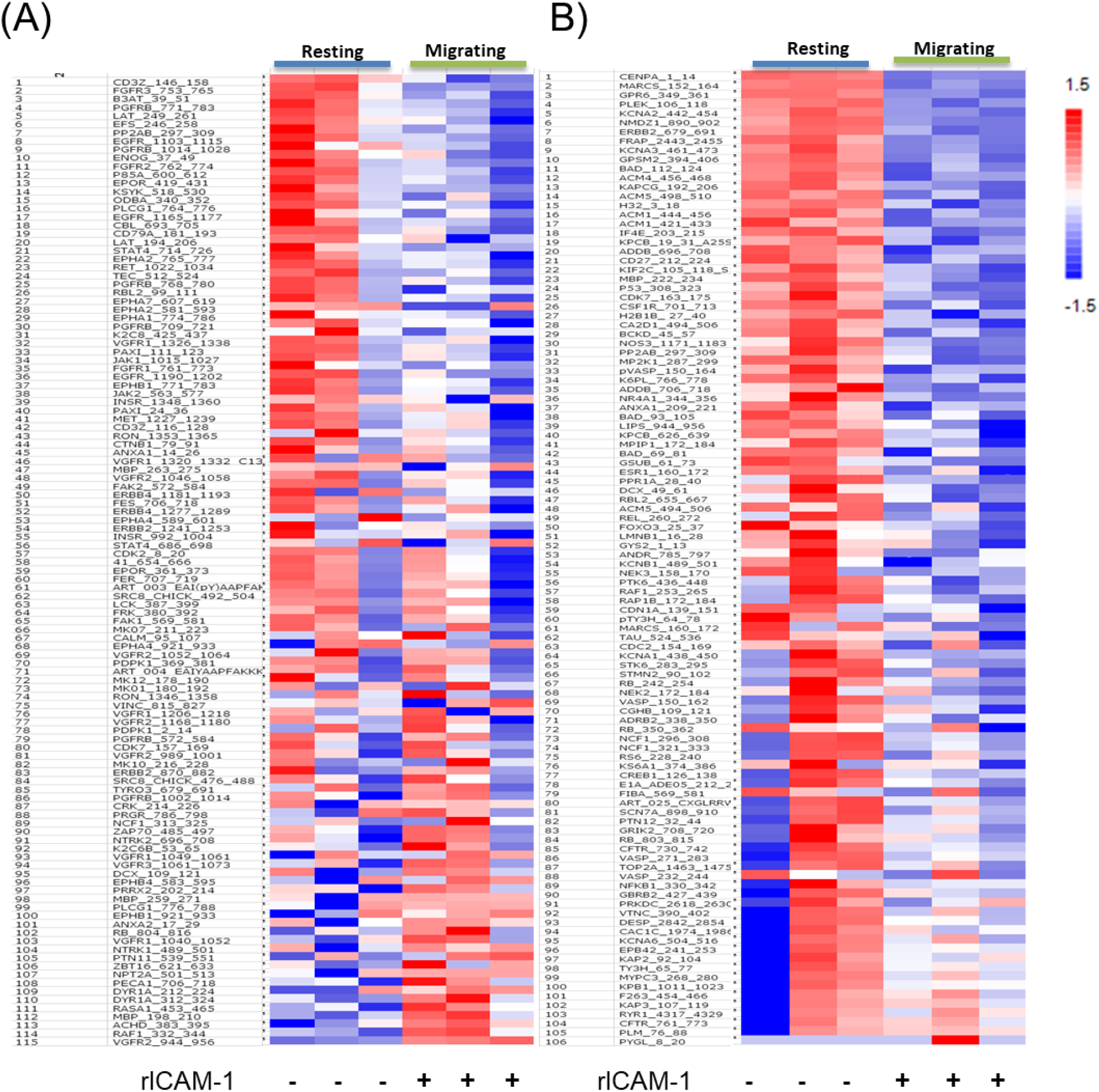
Heat-maps showing phospho-peptide intensities. Human primary T-cells were seeded to migrate on a rICAM-1-coated plate for 30 min and lysed. Cellular lysates were processed for PamChip Peptide array analysis. Heat-maps were plotted with calculated signal values based on intensity calculations of phosphorylated peptides on PTK array (**A**) and STK array (**B**). Peptide Uniprot ID’s along with the range of amino acid sequences employed in the array are indicated on the left side of each panel.

**Figure 4.**
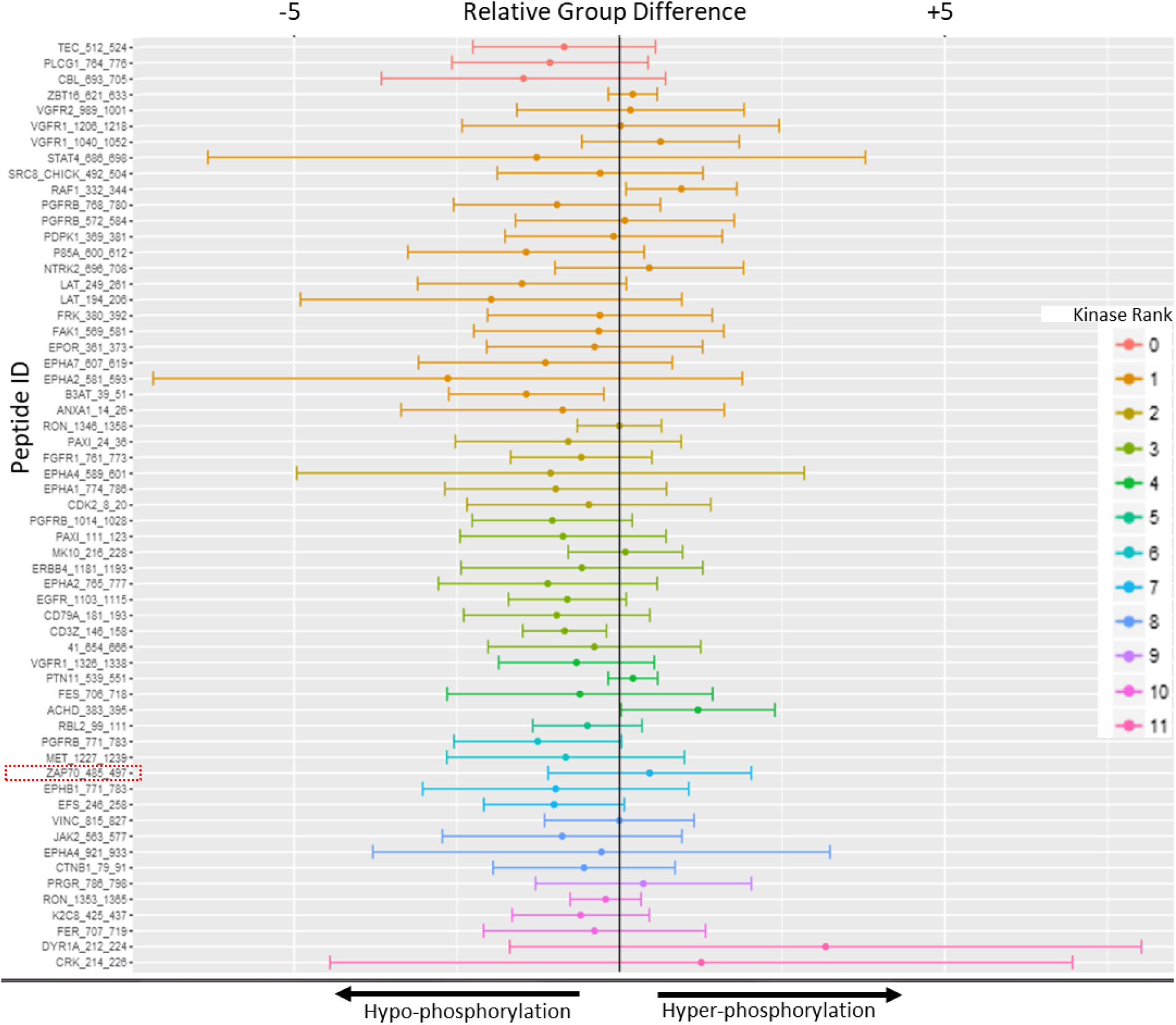
ZAP70-phosphopeptidome analysis in migrating T-cells. A peptide plot with X-axis showing the differences in peptide phosphorylation between LFA-1-stimulated migrating T-cells in comparison to resting controls. The Zap70_485_497 peptide specific phosphorylation shows hyperphosphorylation in migrating T-cells. Upstream kinases analysis of ZAP70-linked-phosphopeptidome indicates a hypophosphorylation of most associated kinases after 30 min of LFA-1 stimulation compared to control. Peptide ranking is based on confidence limit of experimental evidence (*in vitro + in vivo*) and the incidence in all major protein databases - HPRD, PhosphoELM, PhosphositePLUS, Reactome and UNIPROT.

We then performed PamChip® peptide microarray to determine the impact of WFA on downstream cellular kinases in motile T-cells. Kinase specificity scores of WFA-treated T-cells upon stimulation *via* LFA-1/ICAM-1 engagement showed that SYK, ZAP70, AURORA A/B, LCK, and mTOR kinases were among the top kinases constrained by WFA (Fig. 5*A*). In addition, kinase statistical calculations (a measure of kinase activity) predicted TXK, JAK2, FAK, MER, ALK, ZAP70 - PTK’s and PKCε, CAMK4, MAPKAPK2, ERK2 and AKT1 STK’s were inhibited by WFA in LFA-1-stimulated T-cells (Fig. S2). An integrated view of kinase activity modulations by WFA, obtained by upstream kinase analysis, are summarized in the kinase dendrogram (Fig. 5*B*). GeneGo MetaCore® analysis of the kinome datasets designated a role for ZAP70, LCK, LAT, AKT and STAT3 in inhibition of T-cell migration by WFA (Fig. 5*C*). Further validation using Western immunoblot analysis confirmed WFA-mediated inhibition of LFA-1-induced phosphorylation including ZAP70 (pZap70-Tyr319), LAT (pLAT-Tyr191), LCK (pLCK-Tyr505) AKT (pAKT-Ser473), PTEN (pPTEN-Ser380) and AURORA kinase A (pAurora A-Thr288) in migrating T-cells (Fig. 5*D*). At the same time, WFA treatment increased the inactivating phosphorylation of the adaptor protein SLP76 (pSLP76-Ser376) in motile T-cells.

**Figure 5.**
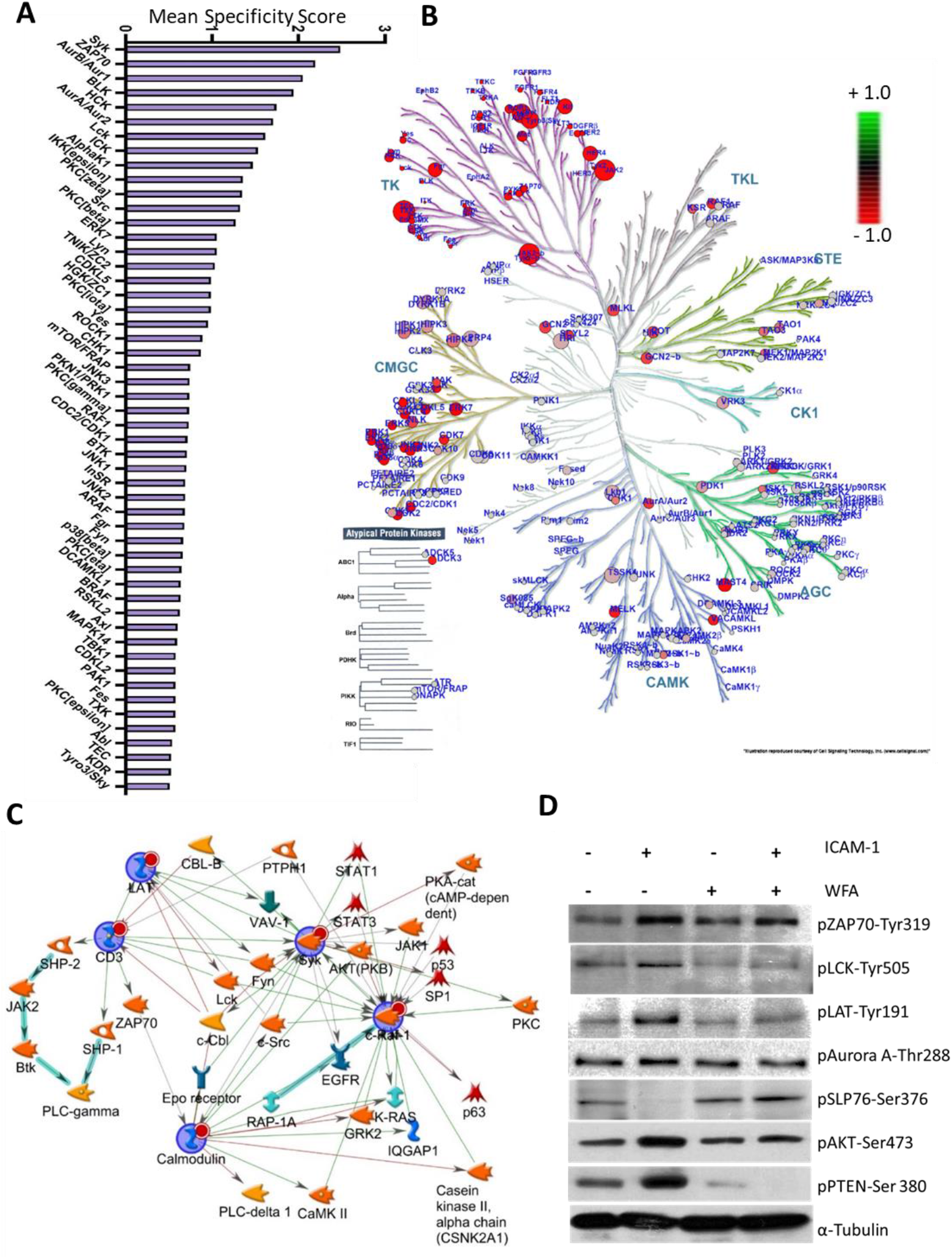
Kinase activity profiling of WFA-treated human T-lymphocytes. (**A**) Mean specificity score calculations signifying the specificity in terms of set of peptides used for the corresponding kinase in WFA-treated migrating T-cells. A higher score indicates higher specificity and less randomness in peptide assignment for the matching kinase. (**B**) Kinase dendrogram complementing the kinome identified in WFA treated LFA-1-stimulated T-cells. Kinases from nearly all groups of the human kinome were detected: CK1, casein kinases; CMGC, CDK/MAPK/GSK3/CLK-family kinases; RCG, receptor guanylate cyclases; STE, sterile homologue kinases; TK, tyrosine kinases; TKL, tyrosine kinase-like kinases; atypical protein kinases. The size of the circle dot indicates the specificity score of the corresponding kinases compared to untreated controls, whereas the colour indicates relative activity (red indicates less active and green hyperactive relative to untreated migrating T-cells control samples). Data represent at least three experiments performed by using T-cells isolated from 3 different donors. (**C**) Network analysis of kinases identified in WFA-treated T-cell kinome. Image generated by GeneGo MetaCore (Clarivate Analytics, USA). (**D**) Western immunoblotting analysis of pZAP70-Tyr319, pLCK-Tyr505, pLAT-Tyr191, pAurora A-Thr288, pSLP76-Ser376, pAKT-Ser473 and pPTEN-Ser380 in WFA-treated T-cells stimulated without or with rICAM-1. The blot was re-probed with anti-α-tubulin as a loading control.

### Molecular modelling predicts WFA binding to ZAP70 at Cysteine 560 and 564 residues

We validated the WFA-ZAP70 interaction, that was identified in the proteomics analysis, by a bait-pulldown approach of ZAP70-WFA-Biotin complex on streptavidin. Western immunoblot probing with anti-ZAP70 showed specific bands associated with ZAP70 both in resting and LFA-1-stimulated T-cells (Fig. 6*A*). Moreover, this interaction was blocked by a competitive thiol donor dithiothreitol (DTT) (Fig. 6*A*). The WFA-ZAP70 interaction was further verified by molecular imaging of WFA-biotin using Alexa Fluor® 555 streptavidin (Fig. 6*B*). Molecular modelling and *in silico* docking predicted hydrogen bonding with amino acid residues L603 and E563 of ZAP70 (Fig. 6*C*). The interaction also included hydrophobic amino acid residues *viz*. P434, V435, S436, C560, C564, P565, P566 and L600. DTT is known to reverse WFA-mediated suppression of kinase activity by blocking alkylation of thiol-sensitive redox pathways (10). Given that DTT-treated human primary T-cells did not show WFA-Biotin and ZAP70 interaction after immunoprecipitation, the most probable thiol donors in the docking complex were either C560, C564 or both residues. As such, the therapeutic efficacy of WFA against different cell types may depend on its promiscuous nucleophilic cysteine reactivity towards multiple hyperactivated cellular kinases.

**Figure 6.**
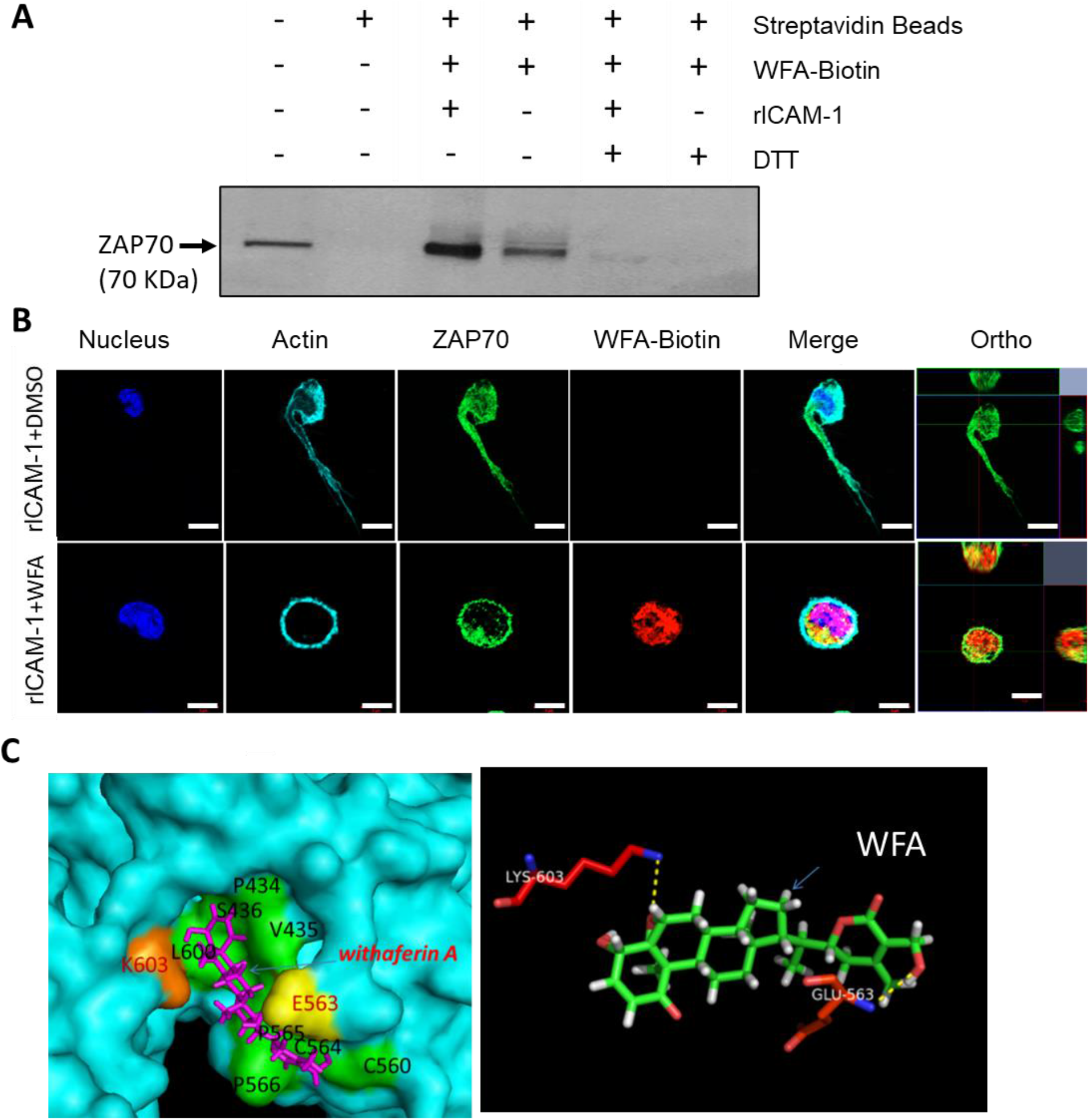
WFA binds to ZAP70 in human T-cells through covalent interactions. (**A**) Human primary T-cells pre-treated with WFA-Biotin for 3 h or DMSO were stimulated to migrate on rICAM-1-coated plates for 30 min and lysed. Cellular lysates were immunoprecipitated with streptavidin-coated agarose beads and then Western immunoblotted for ZAP70. Cellular lysates from DTT-treated cells were evaluated for WFA-Biotin binding as control, whereas T-cell lysates not mixed with WFA-Biotin were used as an in-put control. (**B**) LFA-1-stimulated T-cells either treated with DMSO or WFA-Biotin were immuno-stained with anti-ZAP70/Alexa Fluor® 488 (*green*), streptavidin/Alexa Fluor® 555 (*red*), phalloidin-Alexa Fluor® 647 (actin, *cyan*) and Hoechst (nuclei, *blue*) and then imaged by confocal microscopy. Data represent at least three independent experiments. Scale bar = 5 μM. (**C**) *In silico* analysis of WFA-ZAP70 interaction indicating hydrogen bonding and hydrophobic interactions.

### WFA abrogates LFA-1/ICAM-1 stimulated phosphorylation of MLC-Ser19

We next examined WFA regulation of T-cell cytoskeletal proteins - tubulin and actin. Although the characteristic migratory phenotypes were lost in WFA-treated T-cells, confocal imaging indicated minimal changes in tubulin depolymerization and tubulin bundling architecture in WFA-treated T-cells (Fig. 7*A*). Consistently, the shift in tubulin detyrosination upon WFA treatment compared to untreated migrating control T-cells was limited (Fig. S3). Likewise, there was marginal disparity in nucleation from microtubule organizing centre of acetylated tubulin in WFA-treated T-cells compared to untreated controls (Fig. S3). Furthermore, in microtubule regrowth assays after 15 sec of recovery, we observed the occurrence of microtubule nucleation centres both in WFA-treated and untreated T-cells (Fig. 7*B*), suggesting that tubulin nucleation may not be a prime target for WFA-specific inhibition of T-cell migration. Of note, a reduction in phosphorylated tubulin was observed using p-Tyr272 anti-tubulin antibody (Fig. 7*C*). Disrupted tubulin architecture could be observed upon prolonged incubation of WFA with T-cells (Fig. S4). We argue that WFA could abrogate early phase cytoskeletal dynamics by targeting actin-binding proteins, observed in the WFA-interactome in migrating T-cells. Confocal imaging and Western immunoblotting further confirmed the activity of WFA in depleting pMLC-Ser19 levels in T-cells (Fig. 7*D*, and *E*). Overall, these datasets indicate that WFA abolishes LFA-1 induced phosphorylation of MLC-Ser19 in motile T-cells.

**Figure 7.**
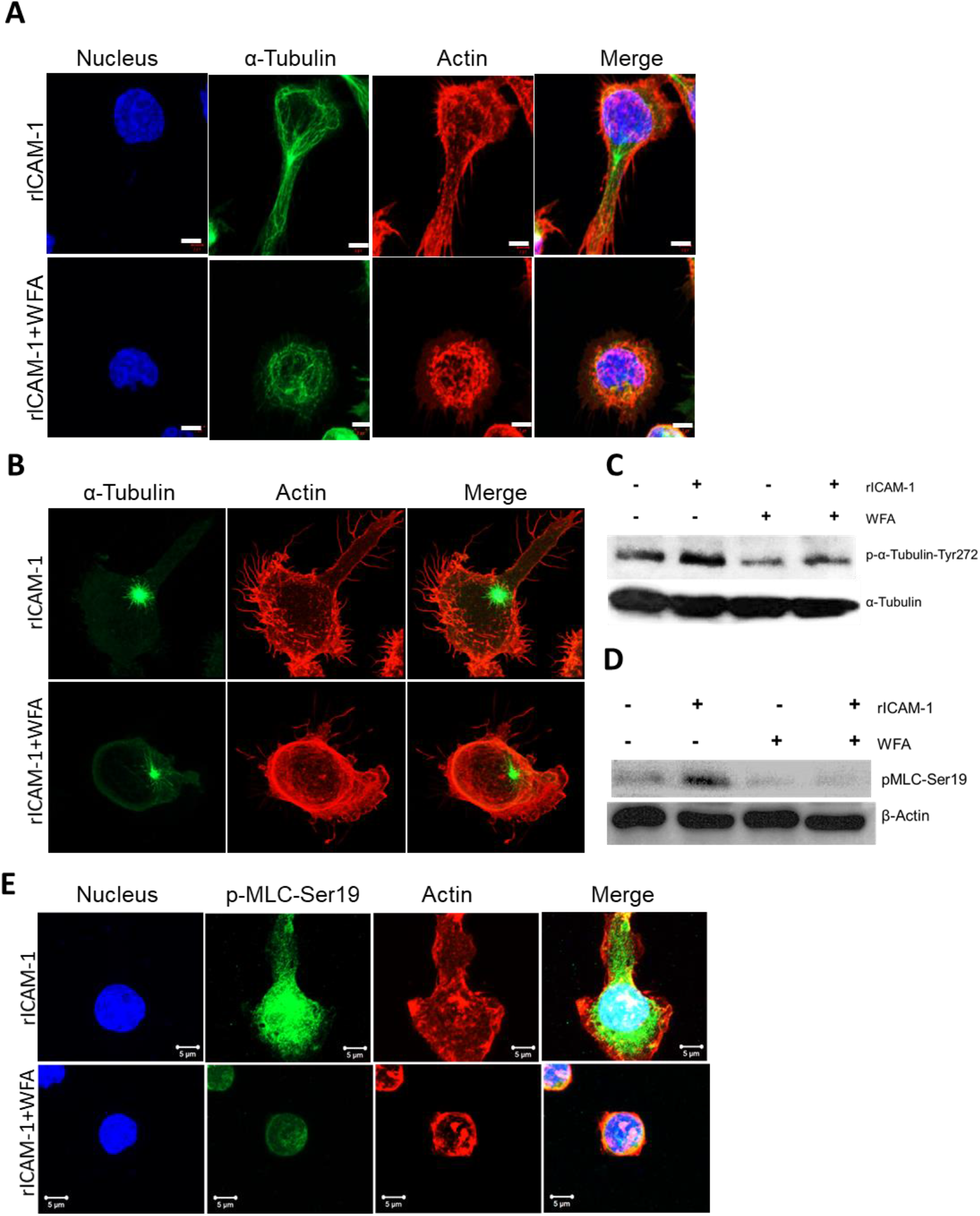
WFA abrogates MLC phosphorylation in motile T-cells. (**A**) Human primary T-cells either treated with 1.25 µM WFA or DMSO were stimulated to migrate on rICAM-1-coated surfaces for 30 min and fixed or lysed. Confocal microscopy images of T-cells stained with anti-α-tubulin-FITC (*green*), Alexa Fluor® 568 Phalloidin (*red*), and Hoechst (nucleus, *blue*). (**B**) Control or WFA-treated T-cells seeded on rICAM-1-coated coverslips were subjected to microtubule depolymerization by incubating cells at 4°C. Microtubule regrowth was analysed by washing cells with prewarm medium and fixing after 15 sec. AiryScan super-resolution microscopy was performed on cells stained with anti-α-tubulin-FITC (*green*) and Alexa Fluor® 568 Phalloidin (*red*) to visualize tubulin regrowth and actin cytoskeleton, respectively. (**C**) Western immunoblots showing expression levels of p-α-tubulin-Tyr272 and α-tubulin (*loading control*) in unstimulated or LFA-1-stimulated T-cells pre-treated with 1.25 μM WFA for 3 h. (**D**) Western immunoblots showing expression levels of p-MLC-Ser19 and β-actin (*loading control*) in unstimulated or LFA-1-stimulated T-cells pre-treated with 1.25 μM WFA for 3 h. (**E**) Confocal microscopy of untreated or WFA-treated LFA-1-stimulated migrating T-cells stained with anti-p-MLC-Ser19/Alexa Fluor® 488 (*green*), Rhodamine Phalloidin® 568 (actin, *red*) and Hoechst (nucleus, *blue*) indicating intense staining of p-MLC-Ser19 at lamellipodial protrusions in migrating T-cells, abrogated by WFA. Scale bar = 5 μM. Data represent at least three independent experiments.

## Discussion

The findings presented herein propose a novel molecular mechanism of WFA action by specific regulation of LFA-1-stimulated signalling in motile T-cells. Our IPA® network analysis of the WFA interactome demonstrated a crucial liaison among ZAP70, NFκB, tumour necrosis factor-α (TNF-α), and chemokine ligand 5 (CCL5) to terminate T-cell migration.

Previous studies have shown that WFA may improve functional recovery in several inflammation model systems, presumably by inhibition of NFκB pathway (11–14). The identification of TNF-α-targeting WFA-interactome associated *via* ZAP70 and NFκB in T-cells could be important in understanding T-cell dysfunctions acquired through prolonged exposure to TNF-α in autoimmune diseases (15). Further examination of WFA interactions with glucose-6-phosphate dehydrogenase, fatty acid synthase, and mTOR identified in the chemo-proteomic analysis will reveal WFA function in redox sensitive immunometabolism and nutrient signaling networks in T-cell migration and inflammation (16).

The WFA-specific phosphopeptidome analyses performed here revealed specific repression of various phospho-Tyr and phospho-Ser/Thr peptides, which is consistent with previous whole cell phosphoproteomics performed in activated T-cells (17). Previously, we and others have demonstrated the importance of ZAP70, PKC isoforms, AKT, TALIN1, and MAPK signalling pathways in LFA-1-induced T-cell migration (18,19). The identification of MET, AKT, ERK2, PKCε and FAK kinases in the LFA-1-stimulated T-cell, emphasizes the importance of PI3K-complex signaling in T-cell extravasation and its inhibition by WFA upon treatment. In fact, various pharmacological inhibitors of PI3K are known to reduce inflammation and mitigation of PI3K signaling is extensively studied in attenuation of autoimmune diseases (20,21).

The current research presents a comprehensive picture of WFA’s activity by elaborating its inhibitory role in T-cell receptor (TCR) related-kinase signalling (*e.g*., ZAP70, LAT, LCK, TXK). Hyperresponsiveness of T-cells to TCR ligation is a hallmark of inflammatory diseases. The role of ZAP70 activation through phosphorylation at various sites in T-cell migration and function has been previously described (5). Our data suggests that the hyper-phosphorylation of ZAP70 to active state in migrating T-cells could be abrogated by WFA. This inhibition of ZAP70 activity by WFA decreased active phosphorylation of LAT-SLP-76 signalosome, implying suppression of downstream TCR-kinase signals. Moreover, we show that WFA interacts with ZAP70 in T-cells that could be abrogated by excess amounts of thiol donor DTT, predicting a favourable WFA binding to cysteines 560 and/or 564 as potential thiol donors in the inhibition of ZAP70 activity. Of note, a C564 mutation in the tyrosine kinase domain of ZAP70 was previously reported in severe combined immunodeficiencies (22).

Translation of T-cell adhesion to cellular migration or function is achieved *via* mobilizing actin linkers (*e.g.*, Talin-1), protein kinase C (PKC) β1- and PKCε-dependent microtubule rearrangements and several protein-protein interactions that facilitate LFA-1 outside-in signaling (6,23,24). The potential of WFA to bind and modulate cytoskeletal elements through actin microfilament aggregation or tubulin depolymerization in other cellular model systems has been described (25–27).

Two populations of microtubules, either dynamic (tyrosinated) or stable (detyrosinated) exist in T-cells and the tyrosinated microtubule fraction increases following stimulation (28). Moreover, a SYK-dependent phosphorylation of tubulin was reported in lymphocyte activation (29,30). However, the effect of LFA-1-stimulation that reduces stable microtubule fraction (by decreasing detyrosinated tubulin) remained relatively similar in WFA treated or untreated motile T-cells. Tubulin regrowth assays showed normal tubulin nucleation even in WFA-treated T-cells. This may be due to the concentration and time of treatment employed in our experiments, which targeted early phase dynamics in T-cell migration. Although total α-tubulin did not show any structural change in confocal imaging (*viz.* tubulin bundling or depolymerization) in the context of early dynamics of T-cell migration, the effect of WFA on tubulin-cytoskeleton appears to be mitigating phosphorylation of α-tubulin (Tyr272).

Kinase signalling induces a coordinated polarization of actin cytoskeleton that is required for T-cell motility (31). The instantaneous intracellular changes arising from LFA-1/ICAM-1 ligation in T-lymphocytes in the early phase lead to ROCK or MLCK kinases dependent acto-myosin contraction, to initiate dynamic reorganization of cytoskeleton (32). Post-translational modification of MLC pivots the migratory apparatus at the leading edge of a migratory T-cell to facilitate cytoskeletal modulations (33,34). Phosphoproteomics studies had previously described p-MLC-Ser19 as the primary site of phosphorylation in active T-cells (35). Therefore, the inhibition of p-MLC-Ser19 by WFA in T-cells is a critical factor in cessation of T-cell migration. The LFA-1/ZAP70 complex is known to recruit other proteins, such as Talin1, ARP-2/3, and WASP to the lamella of polarized T-cell to facilitate actin reorganization (36,37). Interestingly, proteins in the LFA-1/ZAP70 complex that help in actin reorganization were found to be modulated by WFA in migrating T-cells in proteome. However, some of these proteins may well have been pulled-down due to scaffolding activity of cytoskeletal elements, rather than immediate interaction with WFA. A comprehensive mapping of WFA activity in T-cells suggested two modes of actin cytoskeletal regulation in early phase migration. One is by inhibition of cortactin and ARP-2/3 complex and the other by regulating α-actinin (Fig. S5).

A better understanding of cellular signaling dynamics and adaptive mechanisms in T-cells is critical for successful therapeutic strategies against chronic inflammatory or autoimmune diseases. Apart from providing a catalogue of phosphorylation events during LFA-1/ICAM-1 engagement in T-cells, present study demonstrates novel kinases as targets of WFA involved in regulation of human T-cell migration and thus has profound therapeutic implications for inflammatory conditions.

## Experimental procedures

### T-cell culture

Human peripheral blood lymphocyte T-cells were expanded from blood packs obtained from the Health Services Authority of Singapore, as described previously (38). Institutional guidelines were respected in the implementation of experiments and were approved by the Institutional Review Board of Nanyang Technological University, Singapore (IRB-2014-09-007). The human T-cell line HuT78 (ATCC TIB-161TM) was cultured as described previously (39). Briefly, cells were cultured in RPMI-1640 containing 10% foetal bovine serum, 2 mm L-glutamine and Pen-Strep antibiotics (Thermo Fisher Scientific, Singapore) in a humidified chamber at 37°C containing 5% CO_2_.

### High content analysis

A previously optimized high content analysis protocol for T-cell migration and phenotypic quantification was used (40). T-cell migration was induced by incubating cells on immobilized rICAM-1 (Sino Biologicals, China), as described (40). T-cell resting-controls were incubated on poly L-lysine (PLL)-coated plates and cells pre-treated with taxol were used as positive control for migration. WFA (Chromadex, USA) at various concentrations (0.3 μM to 2.5 μM) were used to treat T-cells prior to migratory stimulus. At the end of the experiments, cells were fixed with 3% (*v/v*) formaldehyde in phosphate buffered saline (PBS), fluorescently stained with Alexa Fluor® phalloidin 647 or Rhodamine-Phalloidin (Life Technologies, Singapore) and Hoechst 33258 (Sigma-Aldrich, Singapore). Plates were then scanned (16 randomly selected fields/well at 20X objective) using the IN Cell Analyzer 2200 automated microscope (GE Healthcare, Singapore). T-cell migratory phenotypes and cell shapes were automatically quantified into calculated cell area, nuclear displacement, cell 1/form-factor, and cell numbers. The data were normalised and converted to box and violin plots for better visualization by GraphPad Prism® (Version 8.0, GraphPad Software, USA).

### Trans-well migration assay

Real-time trans-well migration of T-cells was monitored at 37°C using the CIM-Plate 16 and xCELLigence impedance-based detection system (ACEA biosciences, Singapore) to evaluate transmigration of T-cells towards the chemoattractant SDF-1α (Sigma-Aldrich, Singapore) for 16 h, as described (40).

### Cell viability assays

Cell viability was determined using CellTiter 96® Aqueous One solution according to the manufacturer’s instructions (Promega, Singapore).

### Affinity purification and LC-MS/MS analysis

WFA-Biotin was synthesized as previously described (WFA-Biotin; University of Antwerp, Belgium) (41). For identification of endogenous WFA target proteins, both resting and migrating T-cells were incubated for 3 h with WFA-Biotin. Cells were lysed in 0.5 ml lysis buffer and incubated on streptavidin agarose beads (Thermo Fisher, Singapore) overnight and then resolved by SDS-PAGE. Lysates of DMSO-treated T-cells were used as technical controls for mass spectrometry studies.

Tandem mass spectrometry was performed on the *in-gel* digested proteins with trypsin (Promega, Singapore), as described earlier (42). Briefly, peptides were separated using PepMap C18 (3 μm, 100 Å, Thermo Fisher Scientific, MA, USA) and analysed using a Dionex Ultimate 3000 RSLCnano system coupled to a Q Exactive tandem mass spectrometry (Thermo Fisher Scientific, MA, USA). Separation was performed using solvent A (0.1% formic acid) and solvent B (0.1% formic acid in 100% ACN) at flow rate of 300 nL/min with a 60 min gradient. A full MS scan (350– 1,600 m/z range) was acquired at a resolution of 70,000 and a maximum ion accumulation time of 100 ms. The automatic gain control (AGC) settings of the full MS scan and the MS2 scan were 5E6 and 2E5, respectively. An isolation width of 2 m/z was used for MS2. Single and unassigned charged ions were excluded from MS/MS. Raw data files were processed and converted to Mascot generic files format and MS/MS spectra were and submitted for database searching against the Swiss-Prot Human database with Mascot (v2.4.1, Matrix Science). Mascot search results were submitted in HTML format (Online Resource 1). Data generated by Mascot were validated using Scaffold (version 4.10.0, Proteome Software, Portland, OR USA). Protein identifications were accepted if they assigned at least two unique peptides with at least 99% probability. A protein network was generated using Ingenuity Pathway Analysis® (IPA®), by uploading a .csv file containing the list of proteins generated by Scaffold. The network was then improved using the “Build” tool in IPA® to expand the network among the list of proteins identified in the T-cell migration specific WFA-interactome.

### Kinome profiling

T-cell kinome profiling was performed as described previously (43). Briefly, 1 μg T-cell protein lysate from each sample was utilized in the PTK and STK Kinase PamChip® peptide microarrays (PamGene, ‘s-Hertogenbosch, the Netherlands) according to the manufacturer’s protocol in triplicate. Protein lysates of stimulated (*via* LFA-1/ICAM-1 cross-linking) and unstimulated resting or WFA-treated human primary T-cells were loaded to PamChip® arrays containing 115 and 106 immobilized peptides that served as substrates for PTK and STK kinases, respectively. Later, intensities of peptide phosphorylation were monitored by adding a FITC-labelled anti-phospho-tyrosine antibody (PTK array) antibody or a blend of anti-phospho-serine/threonine antibodies in combination with an FITC-labelled secondary antibody. Images were acquired using the CCD camera in the PamStation® 12 and signal quantification on phosphorylated peptides was performed using the Bionavigator software (PamGene International, Den Bosch, Netherlands). Peptide intensities data were log2 transformed and differences in phosphorylation among samples were determined using a non-parametric t-test. Upstream kinase analysis was performed using the PamApp® to identify potential kinases that were hyperphosphorylated or hypophosphorylated compared to respective controls. Kinases were ranked according to their specificity and sensitivity scores (determined by the group and number of peptides phosphorylated). Kinase statistics to measure statistical changes among the experimental groups analyzed, were generated to indicate activity of the identified kinase: a positive score (> 0) reflects kinase hyperactivation, whereas a negative score refers to lower kinase activity in comparison to the control group. The intensity values of peptides showing statistical differences (p-values <0.05) together with the log fold change were exported to GeneGo MetaCore® (Clarivate analytics, USA) for pathway analysis.

### Western immunoblotting

Standard immunoblotting procedures were employed as described previously (40). Equal amounts of T-cell lysates were resolved on SDS-PAGE gels, transferred onto PVDF membranes, incubated with diluted primary antibodies and later with horseradish peroxidise (HRP)-conjugated secondary antibodies to visualize immunoreactive bands by LumiGLO® chemiluminescent detection system (Cell Signalling Technology, Singapore). The images obtained by light sensitive film (Thermo Fisher Scientific, Singapore) or by commercial imaging system (ChemiDoc™, BioRad, Richmond, CA) were analysed by Image J software. All primary antibodies were purchased from Cell Signalling Technology (Singapore) except for rabbit anti-MLC phospho-Ser19 (Sigma Aldrich, Singapore), rabbit monoclonal anti-detyrosinated tubulin and phospho-tubulin Tyr272 (Abcam, Singapore). HRP-conjugated goat antimouse IgG was from Dako (Agilent, Singapore).

### Confocal microscopy

Confocal imaging of T-cells was performed as described previously (40). Briefly, WFA-Biotin or WFA-treated/untreated T-cells were fixed (4% (v/v) formaldehyde) and permeabilized (0.3% Triton X-100 in PBS) to facilitate labelling with appropriate primary and secondary antibodies. Cells were stained with Alexa Fluor® 555 streptavidin (Thermo Fisher Scientific, Singapore) to visualize WFA-biotin, Rhodamine-Phalloidin (Molecular Probes) to visualize the cellular actin and Hoechst 33258 (Sigma-Aldrich) to visualize the nucleus. Confocal imaging was carried out by a laser scanning microscope (Zeiss LSM 800 AiryScan, Carl Zeiss) using a 63X/1.40 N.A. oil immersion objective lens. At least 5 different microscopic fields were analysed for each sample using Zen imaging software (Carl Zeiss).

### Microtubule regrowth assay

Microtubule regrowth experiments were performed as described previously (44). Briefly, microtubule regrowth after tubulin depolymerization in cold shock was analyzed. WFA-treated motile T-cells were washed and fixed after cold shock recovery at 37°C for 15 s and processed for AiryScan confocal microscopy (Zeiss LSM 800 AiryScan, Carl Zeiss).

### WFA-ZAP70 docking

The chemical structure of WFA was retrieved in 2D MDL/SDF format from PubChem database (http://pubchem.ncbi.nlm.nih.gov) before energy minimization procedure in Discovery Studio 2.5 (Dassault Systems BIOVIA). The binding pockets on ZAP70 structure (PDB ID: 1U59) were identified using CASTp server (http://sts.bioe.uic.edu/castp/index.html). Docking was performed using GOLD software (45), as described previously (46). The GOLD score was opted to select the best docked conformations of ZAP-70 in the active site. GOLD parameter file was used to derive empirical constraints used in the fitness function (*viz.*, H-bond energies, torsion potentials, and hydrogen bond directionalities). One complex was selected based on the top GOLD fitness score.

### Statistical analysis

Ordinary one-way ANOVA (for comparison among multiple experimental groups) using GraphPad Prism 4.0 software. For kinome analysis, a non-parametric t-test, two-group comparison, and biological replication comparison tests integrated in the statistical app within BioNavigator software (Pamgene NVA, Netherlands) were utilized. For all data analysis, *p* <0.05 was deemed statistically significant.

## Data availability

All data generated during this study are included in this submitted article and its supplementary information files.

### Funding and additional information

This research was supported, in part, by the Singapore Ministry of Education (MOE) under its MOE Academic Research Fund (AcRF) Tier 2 Grant (MOE2017-T2-2-004), and the National Research Foundation Singapore under its Open Fund Large Collaborative Grant (OFLCG18May-0028) and administered by the Singapore Ministry of Health’s National Medical Research Council (NMRC). W.V.B. acknowledges funding support from the Foundation against cancer (Grant number-7872, Belgium), Hercules foundation (Grant number-AUHA/13/012, Belgium), Research foundation flanders (FWO, grant numbers G059713N/G079614N, Belgium).

## Abbreviations

The abbreviations used are:

DTT: dithiothreitol
ICAM-1: intercellular adhesion receptor-1
IKKβ: I-kappa-B-kinase β
LFA-1: lymphocyte function-associated antigen-1
MAP: mitogen-activated protein
NFκB: nuclear factor kappa B
PBS: phosphate buffered saline
PKC: protein kinase C
PTK: protein tyrosine kinase
rICAM-1: recombinant ICAM-1
SDF-1α: stromal cell-derived factor 1 α
STK: serine/threonine kinase
TCR: T-cell receptor
WFA: Withaferin A
ZAP70: zeta-chain-associated protein kinase 70

